# Exposure to the aryl hydrocarbon receptor agonist dioxin disrupts formation of the muscle, nerves, and vasculature in the developing jaw

**DOI:** 10.1101/2023.06.22.546117

**Authors:** Layra G. Cintrón-Rivera, Nicole Burns, Ratna Patel, Jessica Plavicki

## Abstract

Human exposures to environmental pollutants can disrupt embryonic development and impact juvenile and adult health outcomes by adversely affecting cell and organ function. Notwithstanding, environmental contamination continues to increase because of industrial development, insufficient regulations, and the mobilization of pollutants due to extreme weather events. Dioxins are a class of structurally related persistent organic pollutants that are highly toxic, carcinogenic, and teratogenic. 2,3,7,8-tetrachlorodibenzo-p-dioxin (TCDD) is the most potent dioxin compound and has been shown to induce toxic effects in developing organisms by activating the aryl hydrocarbon receptor (AHR), a ligand activated transcription factor targeted by multiple persistent organic pollutants. Contaminant-induced AHR activation results in malformations in the craniofacial cartilages and neurocranium; however, the mechanisms mediating these phenotypes are not entirely understood. In this study, we utilized the optically transparent zebrafish model to elucidate novel transcriptional and structural targets of embryonic TCDD exposure leading to craniofacial malformations. To this end, we exposed zebrafish embryos at 4 hours post fertilization (hpf) to TCDD and employed a mixed-methods approach utilizing immunohistochemistry staining, transgenic reporter lines, fixed and *in vivo* confocal imaging, and timelapse microscopy to determine the targets mediating TCDD-induced craniofacial phenotypes. Our data shows that embryonic TCDD exposure reduced jaw and pharyngeal arch Sox10+ chondrocytes and Tcf21+ pharyngeal mesoderm progenitors. Exposure to TCDD correspondingly led to a reduction in collagen type II deposition in Sox10+ domains. Embryonic TCDD exposure impaired development of tissues derived from or guided by Tcf21+ progenitors, namely: nerves, muscle, and vasculature. Specifically, TCDD exposure disrupted development of the hyoid and mandibular arch muscles, decreased neural innervation of the jaw, resulted in compression of cranial nerves V and VII, and led to jaw vasculature malformations. Collectively, these findings reveal novel transcriptional and structural targets of TCDD-induced toxicity, showcasing how contaminant exposures lead to congenital craniofacial malformations.

**Highlights:**

- Embryonic TCDD exposure diminishes Sox10+ craniofacial chondrocytes.
- Following TCDD exposure Col2a1 deposition is reduced in Sox10+ domains.
- Exposure to TCDD decreases Tcf21+ progenitors and impairs muscle formation.
- TCDD exposure leads to defects jaw innervation and cranial nerve establishment.
- Early TCDD exposure results in vasculature malformations in the jaw.

**Graphical Abstract:** 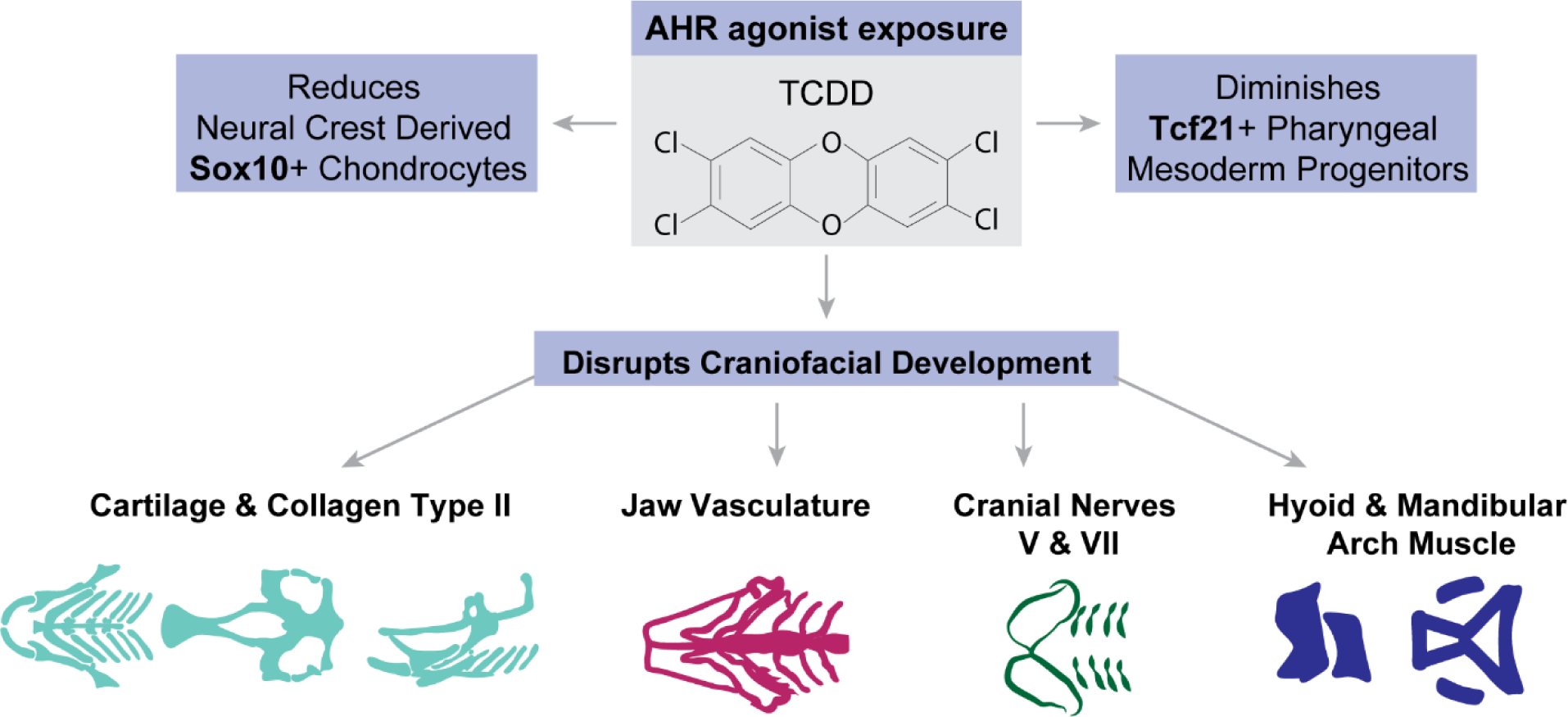

## 1 Introduction

Exposure to environmental contaminants increases the risk of adverse health outcomes; however, the molecular mechanisms, transcriptional targets, and cellular effects of many environmental pollutants remain poorly understood (Briggs, 2003). Of the 30,000 chemicals recognized for common use today only 1% had been evaluated for toxic effects (Haigh, 2004), indicating the extent of pollution-induced health effects may go widely unrecognized. Persistent organic pollutants (POPs) are toxic contaminants that persist in our environment, accumulate in biological systems, and pose a serious risk to human and ecological health. Moreover, due to their chemical stability, legacy contaminants can be mobilized by extreme weather events (Alharbi et al., 2018; Denison & Nagy, 2003). Consequently, emphasis has been placed on understanding the mechanisms of toxicity and the phenotypes induced by POPs. Previous work has found that a subset of POPs, including dioxins, dioxin-like polychlorinated biphenyls (DL-PCBs), dibenzofurans, and polycyclic aromatic hydrocarbons (PAHs) (Alharbi et al., 2018; Denison & Nagy, 2003), primarily induce toxicity through activation of the aryl hydrocarbon receptor (AHR), a ligand activated transcription factor (Carney et al., 2006; Kim et al., 2013; Manzetti & Spoel, 2014; Mimura & Fujii-Kuriyama, 2003; Patrizi & de Cumis, 2018; Sugden et al., 2017; Xu et al., 2015; Zhou et al., 2010)..

Dioxins are a class of structurally related POPs that are highly toxic, widespread, and predominantly generated as byproducts of waste incineration, fossil fuel combustion, and other industrial practices (Dopico & Gómez, 2015; Patrizi & de Cumis, 2018; Rysavy et al., 2013; White & Birnbaum, 2009a). Of the 419 dioxin-related congeners, 2,3,7,8-tetrachlorodibenzo-p-dioxin (TCDD) is the most toxic and potent congener, conferring the highest affinity for and inducing toxicity through activation of AHR (Carney et al., 2006; Kim et al., 2013; Mimura & Fujii-Kuriyama, 2003; Patrizi & de Cumis, 2018; Sugden et al., 2017; Xu et al., 2015). Prior epidemiological and laboratory studies have shown embryonic exposures to TCDD result in congenital craniofacial malformations, among other adverse effects to the nervous, sensory, and cardiovascular systems (Andreasen et al., 2006; Burns et al., 2015; Carney et al., 2006; Guo et al., 2018; Hofsteen et al., 2013; Martin et al., 2022; Nghi et al., 2015; Nghiem et al., 2019; Nishijo et al., 2014; Plavicki et al., 2013; Safe & Luebke, 2016; White & Birnbaum, 2009b; Xiong et al., 2008; Zhang et al., 2018), yet the molecular mechanisms resulting in the observed TCDD-induced and AHR-mediated craniofacial phenotypes have not been fully defined. Therefore, in this study, we utilized the zebrafish (*Danio rerio*) model to systematically characterize the effects of TCDD exposure on embryonic craniofacial development and to interrogate potential transcriptional targets of AHR-mediated toxicity.

Zebrafish provide clear advantages for developmental toxicology research as they develop *ex vivo,* are optically transparent during development, and numerous transgenic tools have been generated that allow researcher to visualize development in real time (Dai et al., 2014; Scholz et al., 2008; Yang et al., 2009). Additionally, 70% of zebrafish genes have a human orthologue (Howe et al., 2013), making zebrafish an excellent model for identifying clinically relevant genes and transcriptional targets of interest. Multiple studies utilizing the zebrafish model have demonstrated that embryonic and larval TCDD exposures result in defective development of the larval jaw cartilages (Burns et al., 2015; Garcia et al., 2018; Xiong et al., 2008). However, the mechanisms mediating these cartilage defects are not fully understood and, while the jaw cartilages have been the focus of prior research examining the effects of embryonic TCDD exposure on craniofacial development, other critical cell types in the jaw have yet to be examined.

Craniofacial development requires complex spatial-temporal regulation of multiple populations of embryonic stem cells, progenitor cells, and differentiated cell types to synchronously establish the tissues required for facial movement and function (Murillo-Rincón & Kaucka, 2020). Among these, the neural crest is a multipotent progenitor population essential for development of the vertebrate head (Santagati & Rijli, 2003). Defects in neural crest expression, establishment, or migration result in craniofacial malformations (Curtin et al., 2011; Quintana et al., 2014; Santagati & Rijli, 2003). SoxE transcription factors, a family of proteins with genetic sequence homology in their high-mobility group (HMG) domains, play a critical role in neural crest and embryonic development. The SoxE family in zebrafish is composed of *sox8*, *sox9a*, *sox9b*, and *sox10*; together these transcription factors are required for embryonic development of the otic vesicle, jaw, heart, fin, gonads, central nervous system, and peripheral nervous system (Hofsteen et al., 2013; She & Yang, 2017; Stolt & Wegner, 2010; Yan et al., 2005). Previous findings indicate TCDD exposure results in downregulation of *sox9b* and leads to jaw cartilage malformations (Xiong et al., 2008). The same study found that morpholino knockdown of *sox9b* was sufficient to recapitulate the cartilage defects observed following exposure to TCDD (Xiong et al., 2008). Moreover, another study found that AHR activation results in expression of a long noncoding RNA, *sox9b long intergenic noncoding RNA* (*slincR*), that represses *sox9b* function and results in abnormal cartilage morphology (Garcia et al., 2018). These results together led to the current mechanistic understanding that TCDD exposure results in AHR mediated downregulation of *sox9b* and subsequent cartilage malformations. Notwithstanding, the role of other SoxE genes in the etiology of toxicant induced craniofacial malformations and the effects of TCDD exposure on other non-cartilaginous tissues remain to be understood.

Therefore, in this study, we exposed 4 hpf embryos to a 1-hour waterborne exposure of 10 parts-per-billion (ppb) TCDD solution and examined the effects early embryonic TCDD exposure on multiple craniofacial tissues and cell types including neural crest derived Sox10+ chondrocytes, collagen type II, Tcf21+ pharyngeal mesoderm progenitors, muscle fibers, nerves, and vasculature. Prior investigations utilizing the zebrafish model have demonstrated that TCDD exposure impacts developmental processes at doses ranging from 1 ppb (Plavicki et al., 2013) to 10 ppb (Souder & Gorelick, 2019). Our lab previously determined that a 1-hour 10 ppb TCDD exposure paradigm results in an internal body burden of 14.7 ±2.04 pg/embryo at 48 hpf and 9.78 ±1.31 pg/embryo at 72 hpf (Kossack et al., 2023). Consequently, we utilized a 10 ppb dosing paradigm in this study to examine the effects of a known body burden on dioxin on craniofacial development. Following embryonic exposure, we used confocal microscopy of fixed and live samples to visualize establishment of critical structures and expression of key transcriptional regulators. Our data reveals previously undescribed transcriptional and structural targets of TCDD in the developing head. Specifically, our data indicate that embryonic TCDD exposure diminishes neural crest derived Sox10+ chondrocytes and collagen type II deposition in Sox10+ domains. We also detected a significant decrease in jaw and pharyngeal Tcf21+ populations, which are derived from Tcf21+ pharyngeal mesoderm progenitors that give rise to muscle and vasculature cells and are required to guide axonal projections. Accordingly, in TCDD exposed fish we observed muscle defects, decreased jaw innervation, and jaw vasculature malformations. Our data indicate that early embryonic exposure to TCDD affects multiple, previously undescribed, transcriptional, and structural targets, offering novel insight into the etiology of congenital craniofacial malformations and the mechanisms of TCDD and AHR-mediated toxicity.

## 2 Methods

### 2.1 Zebrafish Husbandry and Transgenic Lines

All experiments in this study were conducted with zebrafish from the Plavicki Lab Zebrafish Facility. All procedures were covered by protocols approved by the Institutional Animal Care and Use Committee (IACUC) at Brown University in adherence with the National Institute of Health’s “Guide for the Care and Use of Laboratory Animals.” Transgenic zebrafish lines were kept in an aquatic housing system (Aquaneering Inc., San Diego, CA) with automatic pH and conductivity stabilization, centralized filtration, reverse osmosis (RO) water purification, temperature maintenance (28.5 ± 2°C), and ultraviolet (UV) irradiation for disinfection of unwanted microorganisms. Fish were reared and maintained with a 14 hour:10-hour light: dark cycle, according to Westerfield (2000). The Plavicki Lab Zebrafish Facility undertakes routine monitoring for disease including the semiannual quantified Polymerase Chain Reaction (qPCR) panels to detect common fish pathogens.

Adult zebrafish were spawned in 1.7L slope tanks (Techniplast, USA). Fish were situated in tanks 15-18 hours prior to breeding with a transparent divider that separated males and females. The following day the divider was removed within 2-3 hours of the initiation of the light cycle. Embryos were collected over the course of 1 hour and subsequently reared in fresh egg water (60 mg/l Instant Ocean Sea Salts; Aquarium Systems, Mentor, OH) in 100 mm non-treated culture petri dishes (CytoOne, Cat. No. CC7672-3394). Embryonic and larval zebrafish were maintained in temperature regulated incubators with a 14 hour:10-hour light: dark cycle (Powers Scientific Inc., Pipersville, PA). At 24 hpf, embryos were dechorionated manually using Dumont #5XL forceps (Cat. No. 11253-10, Fine Science Tools, Foster City, CA) and moved to fresh egg water containing 0.003 % 1-phenyl-2-thiourea (PTU, Sigma) to prevent pigment deposition. The following transgenic lines were used in this study: *Tg(sox10:RFP)* (Kirby et al., 2006), *TgBAC (cyp1a:NLS:EGFP)* (Kim et al., 2013)*, Tg(isl1:Kaede)* (Barsh et al., 2017)*, Tg(kdrl:GFP)* (Choi et al., 2007)*, TgBAC(tcf21:NLS-EGFP)* (Wang et al., 2011).

### 2.2 TCDD exposure

Embryonic TCDD and dimethyl sulfoxide (DMSO) control exposures were performed in accord with Plavicki et al., (2013) (Plavicki et al., 2013). Within 1 hour of adults spawning, embryos were collected and maintained in a temperature-controlled incubator as described in Methods section 2.1. At 3 hpf fertilized embryos were evaluated for fertilization and the progression of normal development quality. Selected embryos were exposed at 4 hpf to either a 0.1% DMSO vehicle control or to a 10 ppb TCDD dosing solution for 1 hour while gently rocking. Twenty embryos were exposed per 2 mL of dosing solution in 2 mL glass amber vials (Cat. No. 5182-0558, Agilent, Santa Clara, CA). Control and dosing solutions were prepared by adding solution constituents to amber vials already containing 20 embryos and gently rocking. For TCDD exposures, a 10 ppb TCDD (ED-901-B, Cerriliant, St. Louis, MO) dosing solution was prepared by adding 2 mL of egg water and 2 µL of pre-prepared TCDD stock (10ng/mL). Vehicle controls were exposed to a 0.1% DMSO control dosing solution, which was matched to the DMSO concentration in the TCDD dosing solution. Amber vials and caps were wrapped with Parafilm® to limit evaporation. Following exposures, embryos were transferred to 6 well plates and kept in an incubator until screening.

### 2.3 Immunohistochemistry

Zebrafish embryos and larvae were collected at 48, 72, or 96 hpf and fixed in groups of 10-15 fish per 1.5 mL Eppendorf® Safe-Lock microcentrifuge tube (Millipore Sigma, Cat. No. T9661-500EA) in 4% paraformaldehyde (PFA, Sigma-Aldrich, Cat. No. P6148) overnight at 4°C while gently rocking. Post-fixation, samples were washed 5 times with PBS-T (phosphate buffered solution + 0.6% Triton-X 100). Samples were permeabilized with acetone for 5 minutes at −20°C. Post-permeabilization, samples were placed in blocking solution (PBS-T + 4% bovine serum albumin, BSA) overnight while gently rocking at 4°C. After blocking, samples were incubated with 1° antibodies at 4°C for 1 day. Samples next underwent 5 quick washes with PBS-T and then washed overnight in fresh PBS-T at 4°C. Once PBS-T wash solution was removed, the 2° antibody and Hoechst stain (1:10,000) were added to the samples and left to incubate for approximately 18 hours at 4°C. 2° antibody and Hoechst were removed by 5 quick washes performed with PBS-T followed by overnight wash at 4°C in PBS-T. PBS-T solution was fully extracted from the Eppendorf tubes using a Samco™ Fine Tip Transfer Pipette (Millipore Sigma, Cat. No. Z350605-500EA) and VECTASHIELD Mounting Media (Vector Labs, Cat. No. H-1000) was added for clearing and storage. Samples were stored for a minimum of 24 hours in VECTASHIELD prior to confocal imaging. The 1° antibodies used were acetylated-α-tubulin (1:250, mouse monoclonal [IgG2b], Sigma-Aldrich, Cat. No. T7451) and Collagen Type II (II-II6B3) (1:100, mouse monoclonal [IgG1]). II-II6B3 was deposited to the Developmental Studies Hybridoma Bank (DSHB) by Linsenmayer, T.F. (DSHB Hybridoma Product II-II6B3). The 2° antibodies used were Invitrogen™ Alexa Fluor™ Plus 405 Phalloidin (1:100, Fisher Scientific, Cat. No. A30104), Alexa Fluor 647® Goat anti-Mouse IgG2b Cross-Adsorbed Secondary Antibody (1:100, Thermo Fisher, Cat. No. A-21242), and Alexa Fluor® 488 Goat anti-Mouse IgG1 Cross-Adsorbed Secondary Antibody (1:100, ThermoFisher Scientific, Cat. A21121).

### 2.4 Alcian Blue Staining

Zebrafish embryos and larvae were collected and fixed in groups in 4% paraformaldehyde overnight at 4°C while gently rocking. Embryos were then washed in phosphate buffered saline with 0.1% Tween-20 (PBS-Tw). Then embryos were bleached in 30% hydrogen peroxide until transparent. After rinsing in PBS-Tw, embryos were transferred into an 0.1% Alcian blue solution overnight at room temperature while rocking. Embryos were then rinsed for 20 minutes with acidic ethanol (5% hydrochloric acid and 70% ethanol). Finally, embryos were rehydrated through a serial dilution of acidic ethanol and stored in glycerol for future imaging.

### 2.5 Microscopy and Image Analysis

Live embryos and larvae were anesthetized with 0.02% tricaine (MS-222) for embedding and subsequent imaging. Both live and fixed embryos were mounted in 2% low-melting agarose (Fisher Scientific, bp1360-100) in 35 mm glass bottom microwell dishes (MatTek, Part No. P35G- 1.5-14-C). Confocal imaging was performed using a Zeiss LSM 880 confocal with Airyscan Fast. Z-stacks were optimized for sample imaging and spanned approximately 110 μm-230 μm; 10x images collected at 2.60 μm intervals and 40x images collected at 0.49 μm - 0.69 μm intervals. Images were analyzed and prepared for publication using Zeiss Zen Black, Zeiss Zen Blue, Adobe Photoshop, and Adobe Illustrator software. Graphs were generated for data visualization utilizing R studio and GraphPad Prism.

### 2.6 Statistical Analysis

Fish carrying the necessary transgenic constructs were collected at the desired timepoints for analysis. The total n and replicates for control and TCDD-exposed fish has been included for every experiment in its corresponding figure description. A Shapiro Wilk Test was utilized to determine the distribution of quantitative data. Based on data distribution, a parametric t-test with Welch’s correction or a non-parametric t test (Mann Whitney U Test was used to determine significance in GraphPad Prism. To determine differences between categorical data collected from the control and TCDD-exposed groups for vessel analysis a Chi-squared and Fishers exact test were used. Significant differences between groups were determined using Chi-squared and Fishers exact test with a 95% confidence interval in GraphPad Prism. Graphs were produced with R studio and GraphPad Prism. Significance is indicated in each figure as follows: *p < 0.05, **p < 0.01, ***p < 0.001, ****p < 0.0001.

## 3 Results & Discussion

### 3.1 Embryonic TCDD exposure targets cranial neural crest derived Sox10+ chondrocytes

*sox10* is required for early cranial neural crest migration and development of the peripheral nervous system, chondrocytes, inner ear, heart, and melanocytes (Dutton et al., 2009). Previous work from our lab indicates that embryonic TCDD exposure disrupts the development of Sox10+ oligodendrocytes in the brain and Sox10+ inner ear structures (Cintrón-Rivera et al., 2023; Martin et al., 2022); however, it remained unknown whether Sox10+ expression domain in the jaw is also affected by TCDD exposure. Given that prior work in zebrafish has demonstrated TCDD exposure results in jaw cartilage malformations and downregulation of *sox9b* (Xiong et al., 2008). Downregulation of *sox9b* specifically in the jaw was demonstrated after exposure to TCDD at 96 hpf (Xiong et al., 2008). Sox10 is required for chondrocyte establishment and skeletal development (Dutton et al., 2009; Giovannone et al., 2019), we sought to determine if loss of Sox10 contributed to TCDD-induced craniofacial toxicity. We hypothesized that if embryos were exposed to TCDD at 4 hpf, prior to Sox10 neural crest specification and migration (Asad et al., 2016; Lopes et al., 2001), that TCDD exposure would disrupt Sox10+ chondrocyte development.

To characterize the effects of TCDD exposure on craniofacial development and determine if Sox10+ chondrocytes were a potential target of TCDD, we first used confocal microscopy to visualize expression of *cytochrome P450 1A* (*cyp1a*) in the developing zebrafish jaw. Cyp1a is a xenobiotic metabolizing enzyme and a biomarker of AHR activation. Thus, by visualizing expression of Cyp1a, we can identify potential cellular and structural targets of TCDD (Carney et al., 2006; Kim et al., 2013; Mimura & Fujii-Kuriyama, 2003; Xu et al., 2015). To this end, we exposed embryos to TCDD at 4 hpf and then acquired images at 72 hpf of control and TCDD exposed embryos and larvae carrying transgenic reporters for s*ox10* [*Tg(sox10:RFP)*] and *cyp1a* [*TgBAC(cyp1a:NLS:EGFP)*] (Kim et al., 2013; Kirby et al., 2006). TCDD resulted in broad Cyp1a expression in the developing head. Notably, Sox10+ jaw chondrocytes revealed co-expression between Sox10 and Cyp1a, suggesting Sox10+ neural crest derived chondrocytes may be targeted by TCDD (Supp. Fig. 1). Consequently, we sought to determine if TCDD exposure affected establishment of Sox10+ chondrocytes in the jaw and neurocranium, so we analyzed confocal z-series capturing Sox10 expression and quantified the percentage area of the ventral jaw and lateral head covered by Sox10. At 48 hpf, Sox10+ neural crest cells have begun to migrate and establish the presumptive jaw structures (Fig. 1, D) and, by 72 hpf, most of anatomical jaw structures can be distinguished (Fig. 1, A-C, G & J). Our results show that following embryonic TCDD exposure the percentage area of the ventral jaw covered by Sox10+ chondrocytes is significantly reduced at both 48 hpf and 72 hpf (Fig. 1, D-I). Likewise, TCDD exposed fish display a reduction in the percentage area of Sox10+ cells in the lateral head at 72 hpf (Fig. 1, J-L). We were able to visualize the movement of Sox10+ cells through timelapse microscopy, which demonstrated that following embryonic TCDD exposure Sox10+ cells fail to extend into their designated jaw domains when compared to controls. These data suggest TCDD may interfere with the development of Sox10+ neural crest cells, their movement, and/or convergent extension, the processes by which the jaw grows. Furthermore, given that TCDD-induced jaw phenotypes resemble the phenotypes that result from disruptions in planar cell polarity, additional studies should examine if TCDD exposure alters the expression or distribution of critical components of planar cell polarity signaling pathway (Supp. Movie 1)(Kimmel et al., 1998; Le Pabic et al., 2014; Mork et al., 2016).

**Figure 1.**
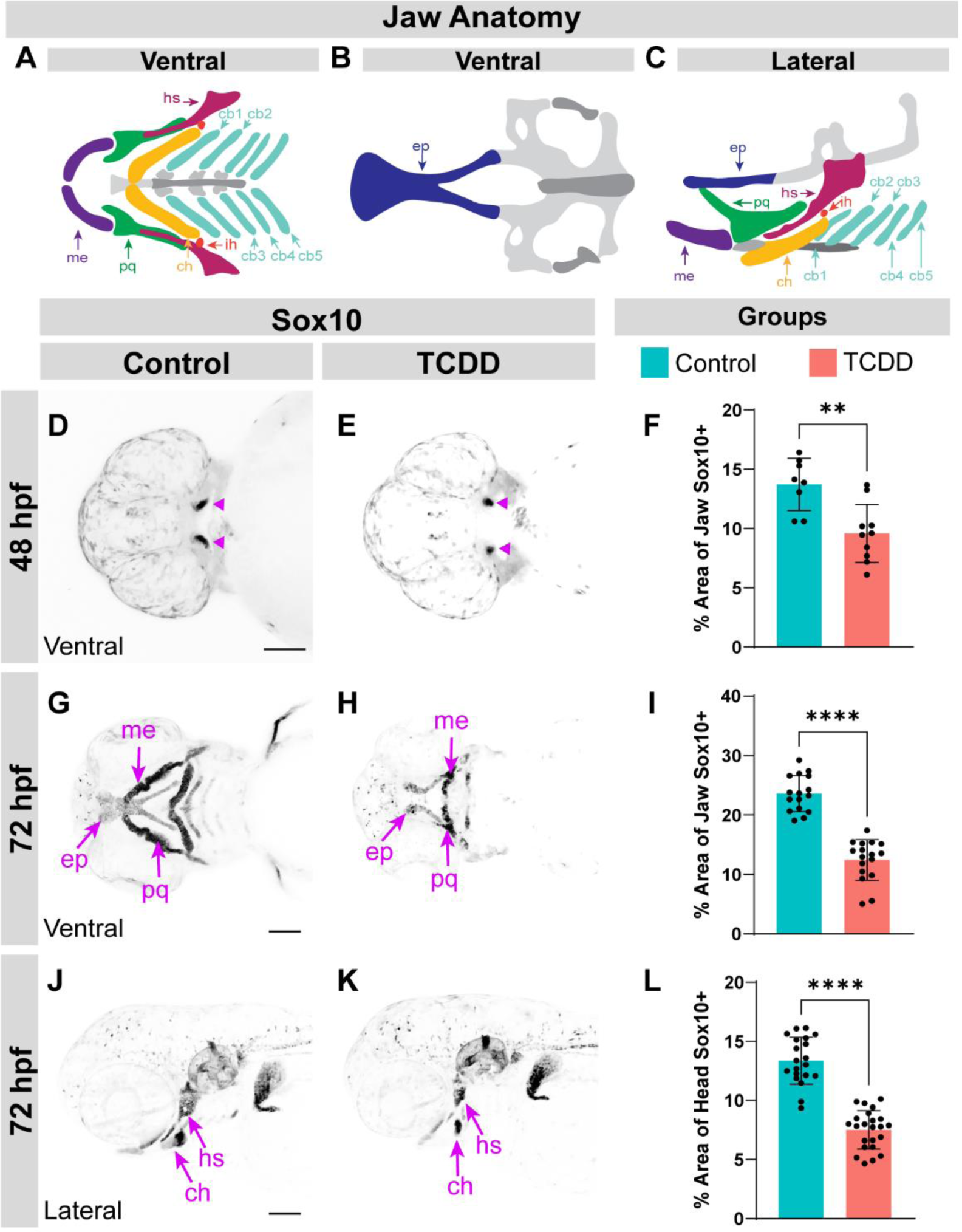
Sox10 expression domain is reduced in the developing jaw and neurocranium following embryonic TCDD exposure. Schematics depicting developed ventral (A-B) and lateral (C) views of wild-type zebrafish jaw anatomy. Schematics were modified and adapted from previously published diagrams (Schilling et al., 1997). Maximum intensity projections of confocal micrographs at 10x magnification capturing control (D, G, J) and TCDD-exposed (E, H, K) embryos and larvae expressing reporters for *sox10* (black) [*Tg(sox10:RFP)*]. Images showcase fish in ventral (D-E, G-H) and lateral (J-K) orientations. Bar graphs indicate percentage area of covered by Sox10+ expression in ventral images at 48 hpf (F) and 72 hpf (I) and in lateral images at 72 hpf (L). Arrowheads indicate primitive Sox10+ jaw structures (D, E). Scale bar in C, G, & J = 100μm. Total n in 48 hpf ventral control group=9 and n in the TCDD-exposed group=10; Total n in 72 hpf ventral control group=22 and n in the TCDD-exposed group=22; Total n in 72 hpf lateral control group=24 and n in the TCDD-exposed group=23. Embryos and larvae were collected across 3 replicates. Abbreviations: hyosymplectic (hs), Meckel’s (me), palatoquadrate (pq), ceratohyal (ch), interhyal (ih), ethmoid plate (ep), ceratobranchial (cb). **p < 0.01, ****p < 0.0001.

TCDD-induced reduction of Sox10+ chondrocytes resulted in overt jaw malformations characterized by absence of multiple, previously overlooked, cartilage structures at 72 hpf as seen through both confocal imaging of the Sox10 transgenic reporter line and Alcian blue staining (Fig. 1 & Supp. Fig. 2). It is worth noting that presence of all jaw structures is required to classify skeletal components of the jaw as these are identified in relation to one another. Therefore, due to the absence of multiple jaw structures in TCDD exposed fish it is difficult to orient positionality in the jaw. Despite these confines, we can report that Meckel’s cartilage, the palatoquadrate, ceratohyal, basihyal, basibranchial, and ceratobranchial structures are consistently absent or smaller in TCDD exposed fish at 72 hpf (Fig. 1, G-H). Moreover, the existing structures seen in ventrally oriented TCDD exposed fish resemble Meckel’s cartilage and the ethmoid plate, but do not have the same characteristics as in controls. Namely, the presumptive ethmoid plate in TCDD exposed embryos is unfused, separated into two distinct structures resembling a cleft palate, and Meckel’s cartilage is in a more posterior coordinate than in controls (Fig. 1, G-H). These findings are consistent with other reports of TCDD-induced craniofacial malformations (Burns et al., 2015; Garcia et al., 2018; Xiong et al., 2008). When observing fish laterally, it also becomes clear that the hyosymplectic and ceratohyal structures are condensed following TCDD exposure (Fig. 1, J-K). The reported reduction in Sox10+ chondrocytes observed at 72 hpf continue at 96 hpf (Supp. Fig. 3). Based on the findings reported in this study, in conjunction with previous data from our lab indicating embryonic TCDD exposure targets Sox10+ oligodendrocytes and the Sox10+ inner ear structures (Cintrón-Rivera et al., 2023; Martin et al., 2022), we conclude that the Sox10+ chondrocytes are a target of early AHR-mediated toxicity and suggest that depletion of Sox10+ chondrocytes may, in part, lead to the previously reported TCDD-induced cartilage malformations.

### 3.2 Collagen type II deposition is reduced in Sox10+ domains following TCDD exposure

Collagens are a broad class of extracellular molecules required throughout the lifespan of an organism for cell adhesion, structural integrity, and cell signaling (Reeck & Oxford, 2022). During early development, a cartilage intermediate made up of chondrocytes embedded in a collagen rich matrix is required for maturation and formation of skeletal structures (Reeck & Oxford, 2022). At least 29 types of collagens have been reported and most have been found in connective tissue, of these, collagen type II (Col2a1) is a major constituent of the chondrocyte secreted extracellular matrix required during development for structural stability and signaling (Reeck & Oxford, 2022; Williams, 2022; Yan et al., 2005). Multiple mammalian studies have found Sox9/SOX9 directly regulates expression of Col2a1 (Bell et al., 1997; Bi, Deng, Zhang, Behringer, & De Crombrugghe, 1999; Zhao, Eberspaecher, Lefebvre, & De Crombrugghe, 1997). Zebrafish studies have confirmed that Sox9a and Sox9b, the zebrafish orthologues of mammalian Sox9/SOX9, directly regulates *col2a1* expression (Chiang et al., 2001; Yan et al., 2005). Avian research has specified that the transcriptional regulation of Col2a1 requires cross regulation of both Sox9 and Sox10 (Suzuki et al., 2006). Previous work conducted in our lab indicated that exposing embryonic zebrafish to TCDD induces inner ear malformations and reduces the expression domain of Col2a1 in the Sox10+ inner ear (Cintrón-Rivera et al., 2023). Based on these prior findings, we surmised that we would see a reduction in chondrocyte associated Col2a1 deposition following the TCDD-induced decrease in Sox10+ craniofacial chondrocytes.

We conducted immunostaining for Col2a1 on control and TCDD exposed fish to determine if TCDD exposure results in diminished Col2a1 expression in the Sox10+ domain of the in developing jaw. At 48hpf, Col2a1 expression was not visible in the developing jaw in neither control nor TCDD exposed (data not shown). Therefore, we focused our efforts on understanding the effects of TCDD exposure at 72 hpf when Sox10+ jaw chondrocytes have begun to stack and form the primitive jaw structures that result in the transcription of and depend on Col2a1. Our results showed that at 72 hpf, following embryonic TCDD exposure, Col2a1 expression is significantly diminished in Sox10+ areas of the jaw, but not in the lens of the eye, when compared to controls (Fig. 2). This robust reduction of Col2a1 deposition in Sox10+ areas, but not in other Col2a1+ domains lacking overlapping or peripheral Sox10 expression, suggests that TCDD exposure may impacts the early transcriptional activity in Sox10+ chondrocytes required for collagen establishment and, ultimately, skeletal formation.

**Figure 2.**
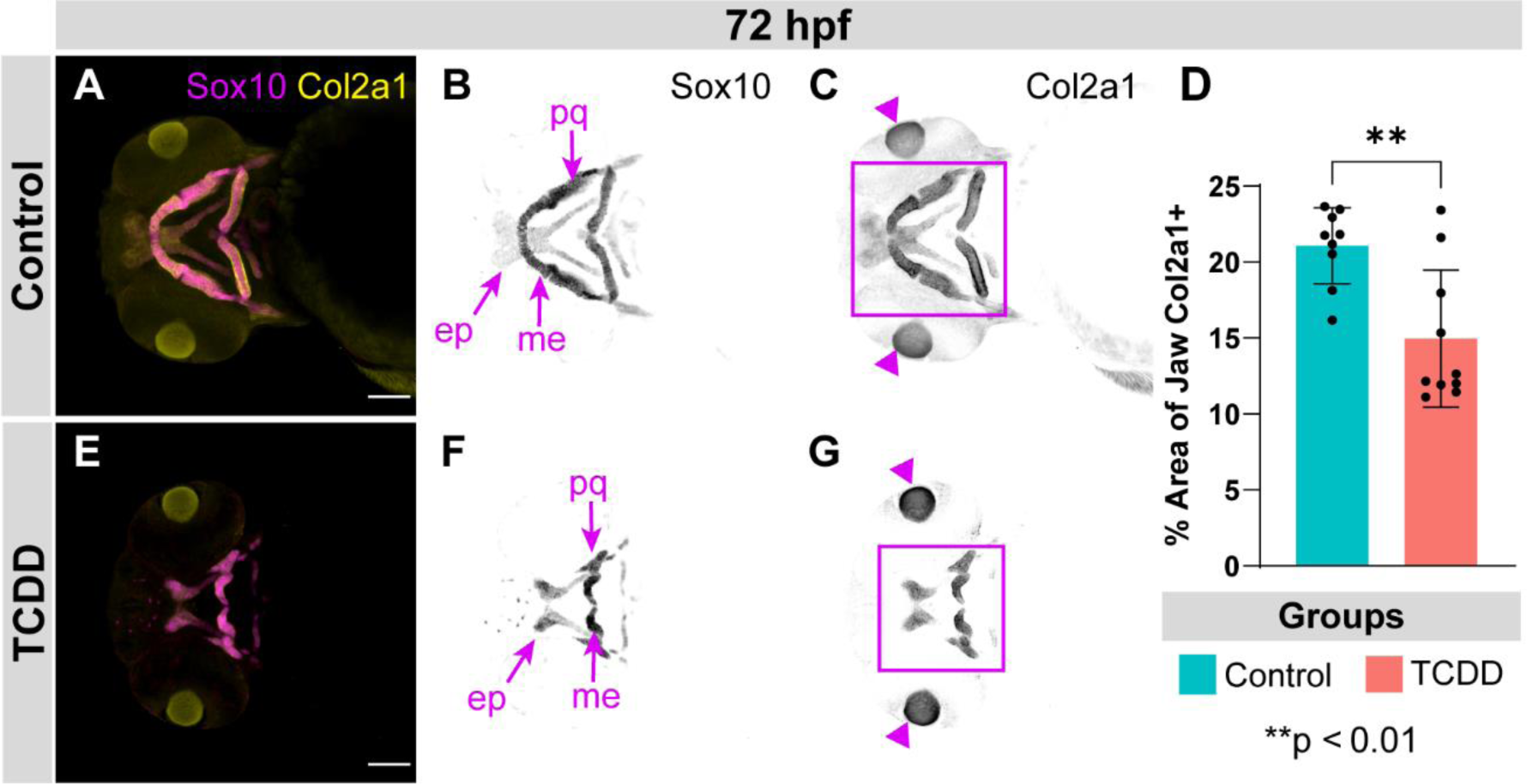
TCDD exposure results in diminished collagen type II deposition. Maximum intensity projections of confocal micrographs at 10x magnification showing control (A-C) and TCDD-exposed (E-G) larvae at 72 hpf. Maximum intensity projections capturing overlapping expression of reporter for *sox10* (magenta) [*Tg(sox10:RFP)*] and, immunostained, Col2a1 (yellow) (A, E). Maximum intensity projections of confocal z-series in black and white showing distinct expression patterns of Sox10 (B, F) and Col2a1(C, G). Square outline designates Col2a1+ jaw domain and arrowheads indicate Col2a1+ domain in the eye (C, G). Bar graph of percentage area of jaw covered by Col2a1+ expression at 72 hpf (D). Scale bar in A & E = 100μm. Total n in control group=9 and n in the TCDD-exposed group=8. Larvae were collected across 3 replicates. Abbreviations: Meckel’s (me), palatoquadrate (pq), ethmoid plate (ep). **p < 0.01.

### 3.3 TCDD exposure diminishes Tcf21+ progenitors and disrupts muscle development

Craniofacial development requires the establishment of and signaling between multiple systems and tissues including cartilage, connective tissue, muscles, nerves, and vasculature. Our initial experiments showed TCDD exposure disrupts development of chondrocytes and their associated collagen type II rich matrix. These results are consistent with past findings while providing new insight into the impact of TCDD exposure on craniofacial development. We next set out to determine if early TCDD exposure affected establishment of craniofacial muscle, another essential tissue required for early developmental signaling, movement, and survival. Tcf21+ progenitors give rise to the majority of craniofacial muscle as well as ventral vasculature (Nagelberg et al., 2015); however, the developmental timeline has not been fully described and Tcf21 expression is seen both within progenitors and differentiated cells. To determine if muscle was a potential target of TCDD toxicity, we utilized the *TgBAC(tcf21:NLS-EGFP)* line and visualized establishment of early muscle progenitors and differentiated cells through confocal microscopy (Wang et al., 2011). Timelapse microscopy experiments demonstrated that Tcf21+ cells fail to migrate to jaw muscle domains following TCDD exposure (Supp. Movie 2). Ventral confocal micrographs of the jaw revealed that TCDD exposure significantly reduced the percent area of the jaw that was Tcf21+ at both 48 and 72 hpf (Fig. 3). Notably, we saw TCDD exposed fish at 72 hpf had diminished or absent expression of Tcf21 at the presumptive intermandibularis anterior, intermandibularis posterior protractor hyoideus, and hyoideus inferior muscle with the remaining Tcf21+ cells largely clustered at the lower jaw area in patterns that did not resemble that of primitive jaw muscle progenitors and cells seen in controls (Fig. 3, F-I). Similarly, lateral images capturing the pharyngeal arch revealed that TCDD exposure diminished the percent area of the pharyngeal arch covered by Tcf21+ cells at both 48 hpf and 72 hpf (Fig. 4). Together, these data indicate that TCDD targets craniofacial Tcf21+ jaw progenitors.

**Figure 3.**
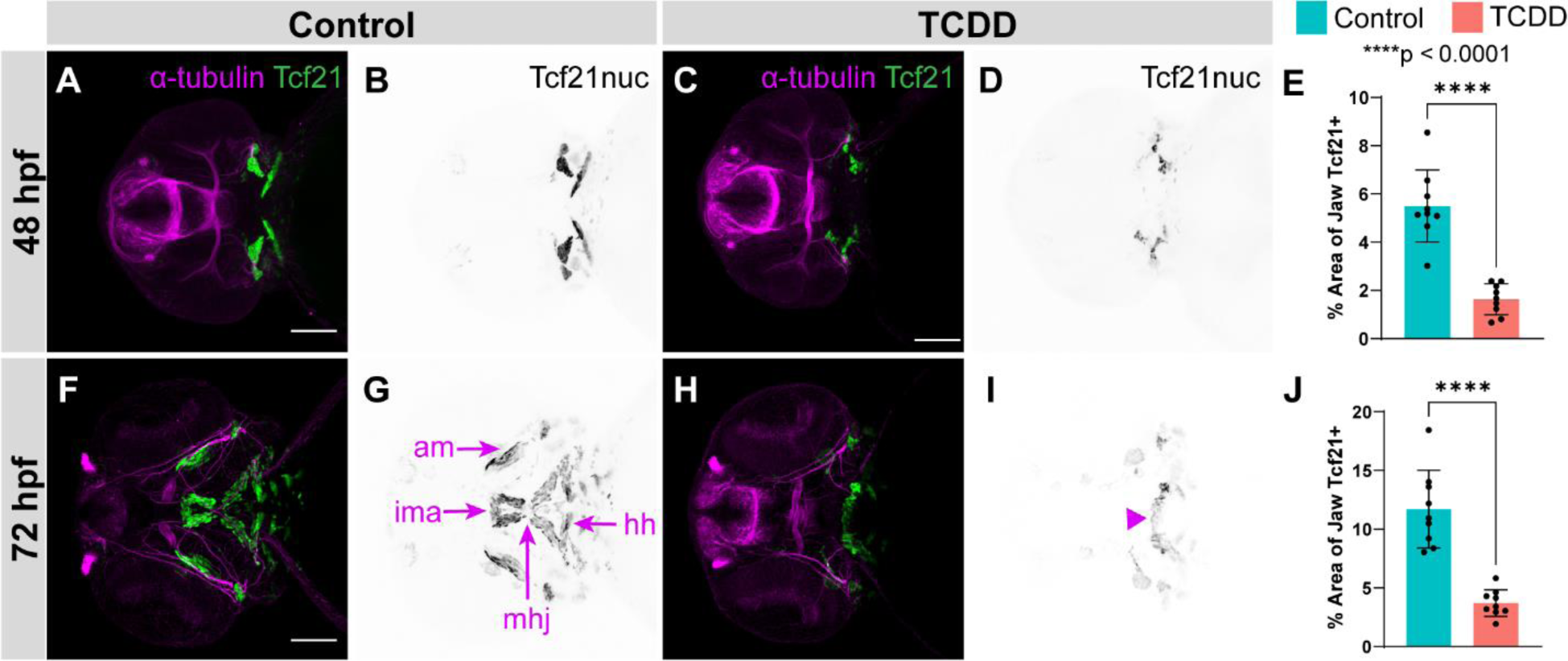
TCDD exposure results in loss of Tcf21 expression in the jaw. Maximum intensity projections of confocal micrographs at 10x magnification capturing control (A-B, F-G) and TCDD-exposed (C-D, H-I) embryos and larvae expressing reporters for Tcf21 (green) [*TgBAC(tcf21:NLS-EGFP)*] and immunostained for acetylated-α-tubulin (magenta) (A, C, F, & H). Maximum intensity projections of confocal z-series in black and white showing distinct expression pattern of Tcf21 (B, D, G, & I). The 48 hpf time points are shown in A-D and the 72 hpf time points are shown in F-I. Bar graph of percentage area of jaw covered by Tcf21+ expression at 48 hpf (E) and 72 hpf (J). Arrowhead highlights Tcf21+ cells clustered at the posterior of the developing head in TCDD exposed fish at 72 hpf (I). Scale bar in A & F = 100μm. Total n in 48 hpf control group=9 and n in the TCDD-exposed group=10; Total n in 72 hpf control group=9 and n in the TCDD-exposed group=9. Embryos and larvae were collected across 3 replicates. Abbreviations: intermandibularis anterior (ima), abductor mandibularis (am), hyohyal (hh), mandibulohyoid junction (mhj). ****p < 0.0001.

**Figure 4.**
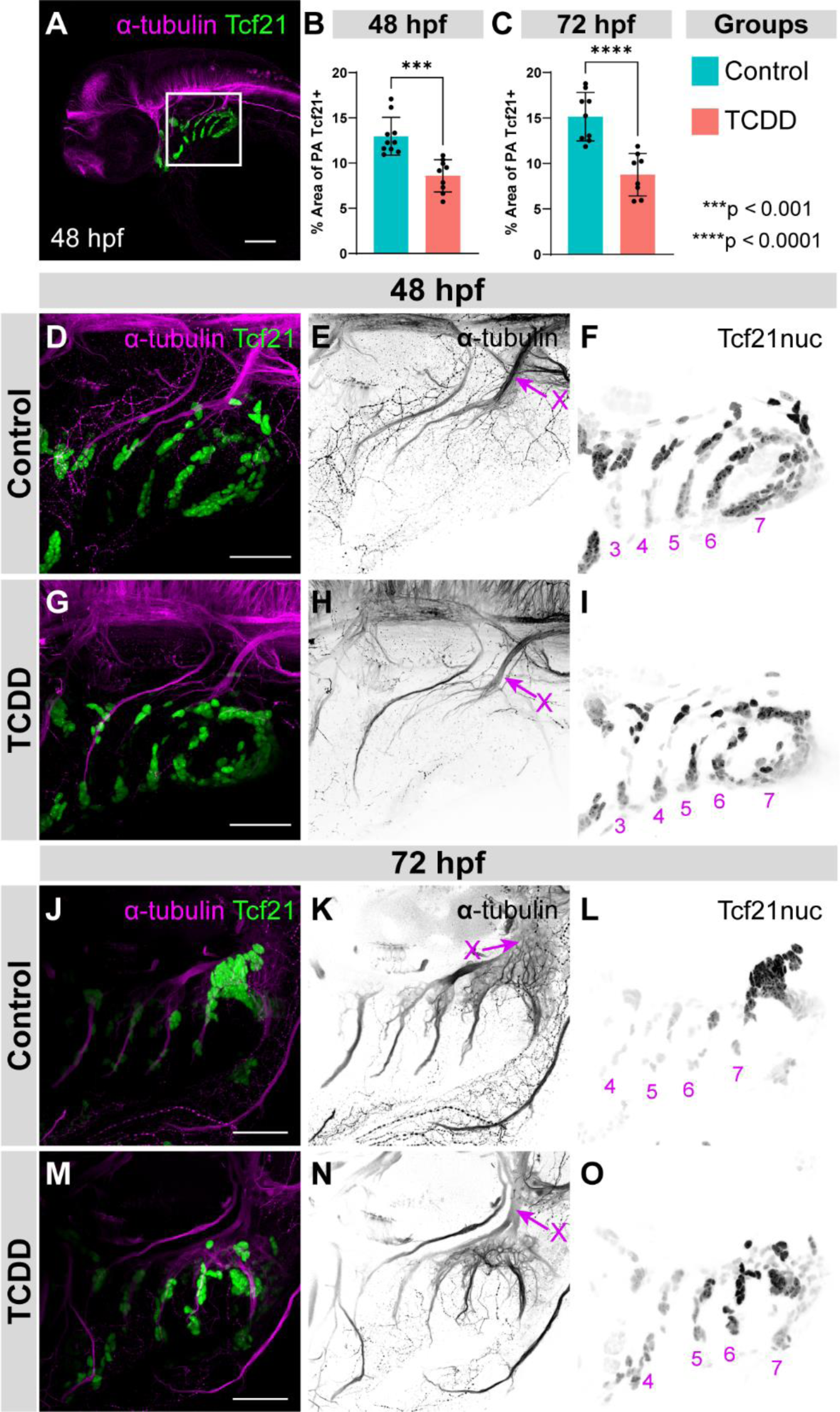
Tcf21 expression in the pharyngeal arch is reduced. Maximum intensity projections of confocal micrographs at 10x (A) and 40x (D-O) magnification capturing control (A, D-F, J-L) and TCDD-exposed (G-I, M-O) embryos and larvae expressing reporters for *tcf21* (green) [*TgBAC(tcf21:NLS-EGFP)*] and immunostained for acetylated-α-tubulin (magenta) (A, D, G, J, & M). White square in A indicates pharyngeal arch region. Maximum intensity projections of confocal z-series in black and white showing distinct expression pattern of acetylated-α-tubulin (E, H, K, & N) and Tcf21 (F, I, L, & O). The 48 hpf time points are shown in A & D-I and the 72 hpf time points are shown in J-O. Bar graphs of percentage area of pharyngeal arch covered by Tcf21+ expression at 48 hpf (B) and 72 hpf (C). PA3-7 are labeled at 48 hpf (F, I) and PA4-7 at 72hpf (L,O). Scale bar in A = 100μm. Scale bar in D, G, J, & M = 50μm. Total n in 48 hpf control group=10 and n in the TCDD-exposed group=9; Total n in 72 hpf control group=10 and n in the TCDD-exposed group=9. Embryos and larvae were collected across 3 replicates. Abbreviations: Vagus Nerve (X). ***p < 0.001, ****p < 0.0001.

Previous work looking at a zebrafish *tcf21* mutant found that deletion of *tcf21* leads to absence or severe reduction of all muscles of the head (Nagelberg et al., 2015). Similarly, mouse studies have also implicated Tcf21 as an essential marker of the pharyngeal mesoderm which derives into craniofacial muscle (Harel et al., 2012). These studies in combination with our findings that TCDD reduces Tcf21+ progenitors and cells led us to postulate that early exposures may, in due course, affect muscle establishment. Therefore, we sought to determine if the significant loss of Tcf21+ progenitor populations was followed by defects in craniofacial muscle formation. To this end, we utilized phalloidin staining, which marks actin filaments to allow for visualization of muscle through confocal microscopy (Higashijima et al., 2000).Our results indicate that TCDD exposure impairs establishment of muscle fibers of the hyoid and mandibular arch muscles. Specifically, the intermandibularis anterior, protractor hyoideus, and hyoideus inferior muscle in the jaw, and the levator arcus palatini, abductor hyomandibulae, dilator operculi, and abductor operculi were repeatedly affected (Fig 5). Our results showed that exposure to TCDD affects muscle filament establishment to varying degrees of severity (Fig 5). Moreover, muscle fiber malformations in TCDD exposed fish do not resolve by 96 hpf, suggesting that defects persist through development and are not the result of a developmental delay (Supp. Fig. 3).

**Figure 5.**
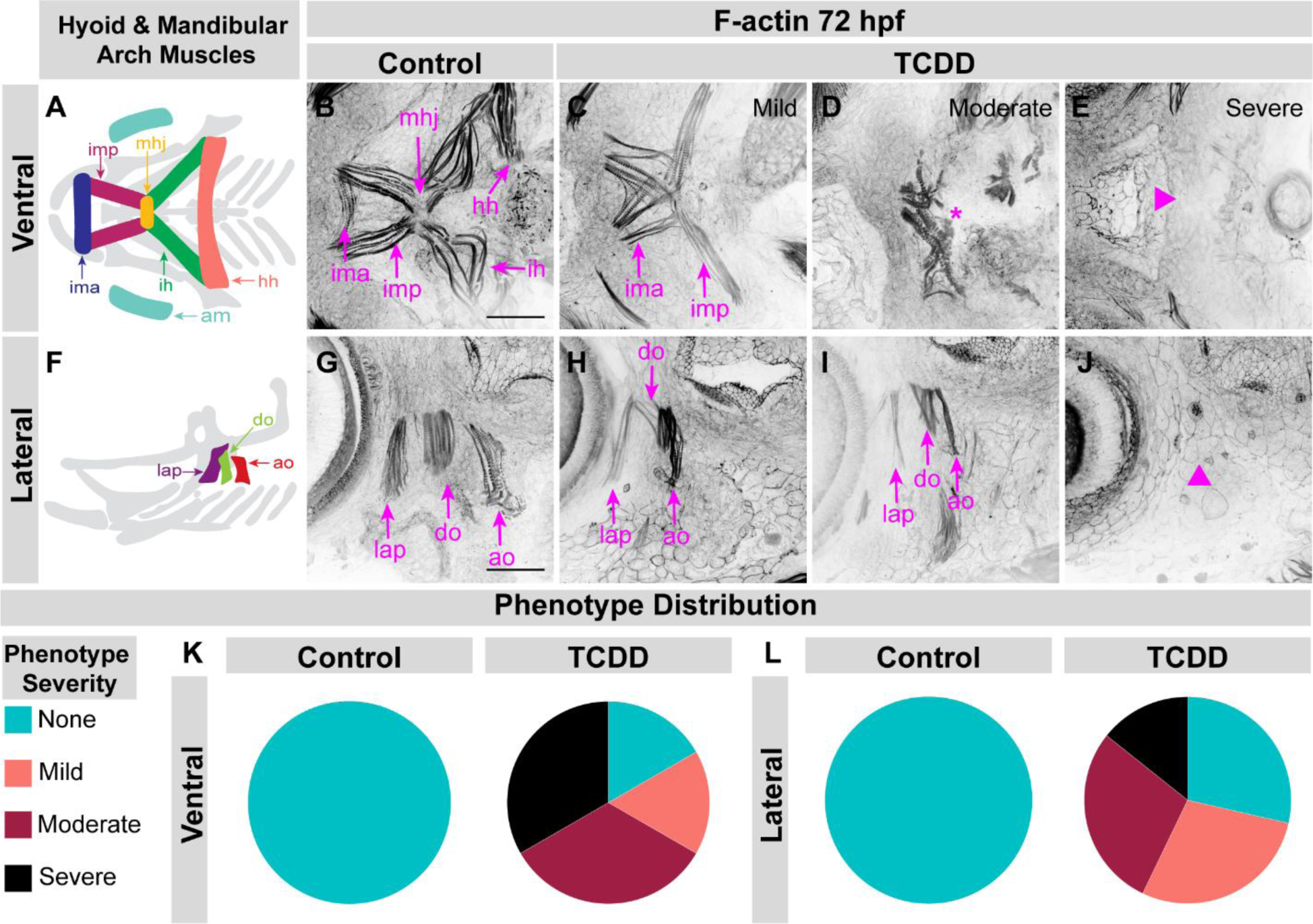
TCDD exposure disrupts muscle development. Schematics illustrating developed ventral (A) and lateral (F) view of wild-type zebrafish hyoid and mandibular arch muscle anatomy (cartilage anatomy shown in grey). Maximum intensity projections of confocal micrographs at 40x magnification showing control (B & G) and TCDD-exposed (C-E, H-J) larvae at 72 hpf stained for F-actin (magenta) to visualize muscle. Asterisk indicates indistinguishable structure of muscle fibers (D). Arrowheads highlight absence of muscle by pointing to area where muscle structures should form (E, J). Pie chart displaying quantification of phenotype distribution for ventral (K) and lateral (L) muscles. Scale bar in B & G = 50μm. Total n in ventral control group=11 and n in the TCDD-exposed group=11; Total n in lateral control group=11 and n in the TCDD-exposed group=14. Larvae were collected across 3 replicates. Abbreviations: abductor mandibularis (am), intermandibularis anterior (ima), intermandibularis posterior (imp), mandibulohyoid junction (mhj), interhyal (ih), hyohyal (hh), levator arcus palatini (lap), dilator operculi (do), and abductor operculi (ao).

To our knowledge, no previous studies had determined the effect of embryonic TCDD exposure on zebrafish craniofacial muscle development until this work. Embryonic TCDD exposure had previously been found to eliminate Tcf21 expression in the zebrafish proepicardium, resulting in a loss of Tcf21+ epicardial cells at 120 hpf (Plavicki et al., 2013). The effects of TCDD exposure on Tcf21+ progenitors in other developmental contexts has not been evaluated in the jaw until this study. Previous work had been done to determine the effects of TCDD exposure on the development of muscle in the red seabream (*Pagrus major*) model (Iida et al., 2013). This study found that an 80 minute exposure to 0.4 ug/L TCDD at 10 hpf did not result in craniofacial muscle development defects as seen through immunohistochemistry staining with pan-myosin heavy chain (pan-MyHC) antibody MF20 when evaluated at 120 hpf (Iida et al., 2013); however, these exposure paradigms significantly differed from that of our study and the lower magnifications used to analyze MF20 expression may not have been sufficient to capture nuanced damage to the developing muscle. In summary, the observed reduction of early Tcf21+ progenitors and differentiated Tcf21+ cells in the jaw and pharyngeal arch, in combination with the muscle fiber malformations of the hyoid and mandibular arch muscles, demonstrate that Tcf21+ pharyngeal mesoderm progenitors, differentiated Tcf21+ cells, and their associated muscle structures are novel targets of TCDD-induced embryonic toxicity.

### 3.4 Jaw innervation is perturbed after TCDD exposure

Cranial nerves are responsible for innervating the muscles of the eye, head, and neck (Guthrie, 2007). Accordingly, the cranial nerves are essential across species for basic functions required for survival such as facial movements, feeding, and, when applicable, vocalized communication (Guthrie, 2007). The branchiomeric nerves (V, VII, IX, and X), a subset of cranial nerves, are the motor and sensory nerves responsible for innervating tissues derived from the pharyngeal arches (Higashijima et al., 2000). Humans require the trigeminal motor nerve (cranial nerve V) to innervate the muscles required for mastication (Chandrasekhar, 2004) and the facial motor nerve (cranial nerve VII) to innervate the muscles required for facial movements and expressions, as well as muscles within the middle ear and neck (Chandrasekhar, 2004). Isl1 is a marker, found in zebrafish and mammalian models alike, of differentiated cranial motor neurons which are a subset of neurons that make up the cranial nerves (Barsh et al., 2017; Higashijima et al., 2000; Uemura et al., 2005). Zebrafish studies have shown Tcf21+ derived muscles signal and guide Isl1+ axon projections of the branchiomeric nerves to innervate the jaw and pharyngeal arch (Barsh et al., 2017).

Given the TCDD-induced decrease in Tcf21+ progenitors and impaired muscle development phenotypes reported earlier in this study, we hypothesized early TCDD exposure would adversely affect jaw innervation. Thus, we employed acetylated-α-tubulin immunostaining and utilized the *Tg(isl1:Kaede)* line to visualize development of axonal projections and cranial motor nerves, respectively (Barsh et al., 2017). Our data indicate that the proportional percent area of the ventral head that is covered by acetylated-α-tubulin is significantly reduced in TCDD exposed fish at 72 hpf, but not at 48 hpf (Fig. 6, E, & Supp. Fig. 4). In contrast, the percent area of the lateral head covered by acetylated-α-tubulin at 72 hpf was not significantly reduced (Supp. Fig. 5), suggesting TCDD exposure results in jaw specific innervation deficiencies. Based on these results, we opted to assess the effects of TCDD on the trigeminal nerve (V) and facial motor nerve (VII), the cranial nerves that innervate the ventral jaw, which are visible when imaging the *isl1* reporter line [*Tg(Isl1:Kaede)*] at 72 hpf (Barsh et al., 2017). Our images capturing *isl1* expression show that following TCDD exposure both the trigeminal nerve (V) and the facial motor nerve (VII) consistently develop with a compressed phenotype (Fig 6, D & I). Specifically, following TCDD exposure the convergence of nerves at the midline, where the jaw muscles intersect at the mandibulohyoid junction, is malformed, indicating a potential branching or targeting defect, which may result from the observed reduction in Tcf21+ cells. Together, these data demonstrate TCDD exposure disrupts development of cranial nerves V and Vll, reduces overall neural innervation of the ventral head, and, ultimately, impairs innervation of jaw, suggesting that TCDD-induced depletion of Tcf21 cues coupled with reduced craniofacial structures may play a role in toxicant-induced innervation phenotypes.

**Figure 6.**
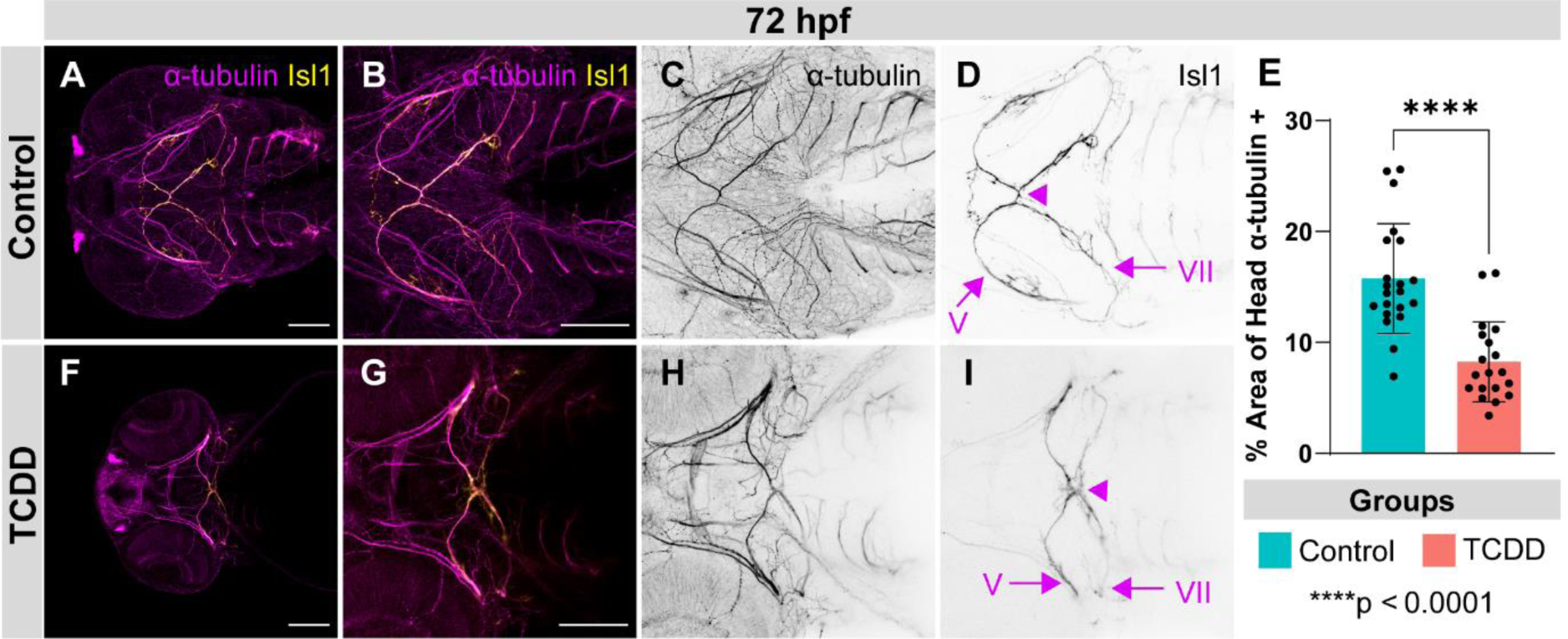
Jaw innervation is disrupted following TCDD exposure. Maximum intensity projections of confocal micrographs at 10x (A, F) and 20x (B-D, G-I) magnification capturing control (A-D) and TCDD-exposed (F-I) larvae at 72 hpf expressing reporters *isl1* (yellow) [*Tg(isl1:Kaede)*] and immunostained for acetylated-α-tubulin (magenta) (A-B, F-G). Maximum intensity projections of confocal z-series in black and white showing distinct expression pattern of acetylated-α-tubulin (C, H) and Isl1 (D, I). Arrowhead indicates the junction of trigeminal motor nerve (V) and facial motor nerve (VII) (D, I). Bar graph of percentage area of head covered by acetylated-α-tubulin expression at 72 hpf (E). Scale bar in A, B, F, & G = 100μm. For acetylated-α-tubulin quantification total n control group=22 and n in the TCDD-exposed group=23. For *isl1* visualization total n control group=10 and n in the TCDD-exposed group=10. Larvae were collected across 3 replicates. Abbreviations: trigeminal motor nerve (V) and facial motor nerve (VII). ****p < 0.0001.

Prior animal studies have found genetic knockouts that lead to abnormal development of the development of both the trigeminal nerve (V) and facial motor nerve (VII). The *Neuropilin2* knockout mouse model results in both cranial nerve V and VII being defasciculated (Chen et al., 2000; Giger et al., 2000) and, in the *Neuropilin2* and *Sema3A* double knockout mouse, the defasciculated phenotype is exacerbated (Chen et al., 2000; Giger et al., 2000). Additionally, mutations in HOXB1 also affects both cranial nerves V and VII and, ultimately, results in facial paralysis (Guthrie, 2007). Therefore, these genes with established roles in both cranial nerve V and VII should be examined in future studies assessing effects of TCDD and AHR-mediated toxicity on cranial nerve development.

### 3.5 Embryonic TCDD exposure leads to jaw vasculature malformations

Craniofacial muscles and jaw vasculature are derived from Tcf21+ pharyngeal mesoderm progenitors (Nagelberg et al., 2015). In this study, we found that TCDD exposure leads to a significant decrease of the Tcf21+ population of cells in the jaw and pharyngeal arches at 48 and 72 hpf followed by complete absence or robust reduction of craniofacial muscle fibers and a reduction in ventral innervation at 72 hpf. Since Tcf21+ progenitors give rise to both muscle and jaw vasculature, we sought to determine if vessels of the developing jaw also failed to form following the embryonic TCDD-induced reduction of Tcf21+ progenitors and differentiated Tcf21+ cells. A preceding study had found that a exposure to TCDD at 24 hpf impaired lower jaw development, as depicted through Alcian blue staining of the cartilage, in a concentration dependent manner and, in due course, led to circulation failure at 72 hpf and altered heart rate at 96 hpf (Teraoka et al., 2002). The same study found that TCDD had vessel specific blood flow effects, that is, TCDD reduced blood flow at the intersegmental vessels, dorsal aorta and posterior cardinal vein while increasing blood flow at the hypobranchial artery (Teraoka et al., 2002). Nevertheless, these aforementioned results were based on a TCDD exposures performed at 24 hpf (Teraoka et al., 2002), after Tcf21+ progenitors have been specified at approximately 15 hpf (Nagelberg et al., 2015). Thus, we hypothesized that a 4 hpf exposure, prior to Tcf21+ specification, would result in more robust vasculature malformations. Accordingly, our results demonstrate that early TCDD exposure impairs development of multiple, previously unspecified, ventral vessels including establishment of the opercular artery and the ventral aorta (Fig 7). 100% of TCDD exposed fish displayed complete absence of the opercular artery and, by consequence of this absence, defects in the vascular connection from ventral jaw to the hypobranchial artery (Fig 7, F). Furthermore, the ventral aorta was absent and thereby lacking adequate connection to other vessels in 100% of exposed fish (Fig 7, F). Collectively, our data indicate that embryonic TCDD exposure leads to a reduction of Tcf21+ progenitors and results in subsequent muscle, nerve, and vasculature defects.

**Figure 7.**
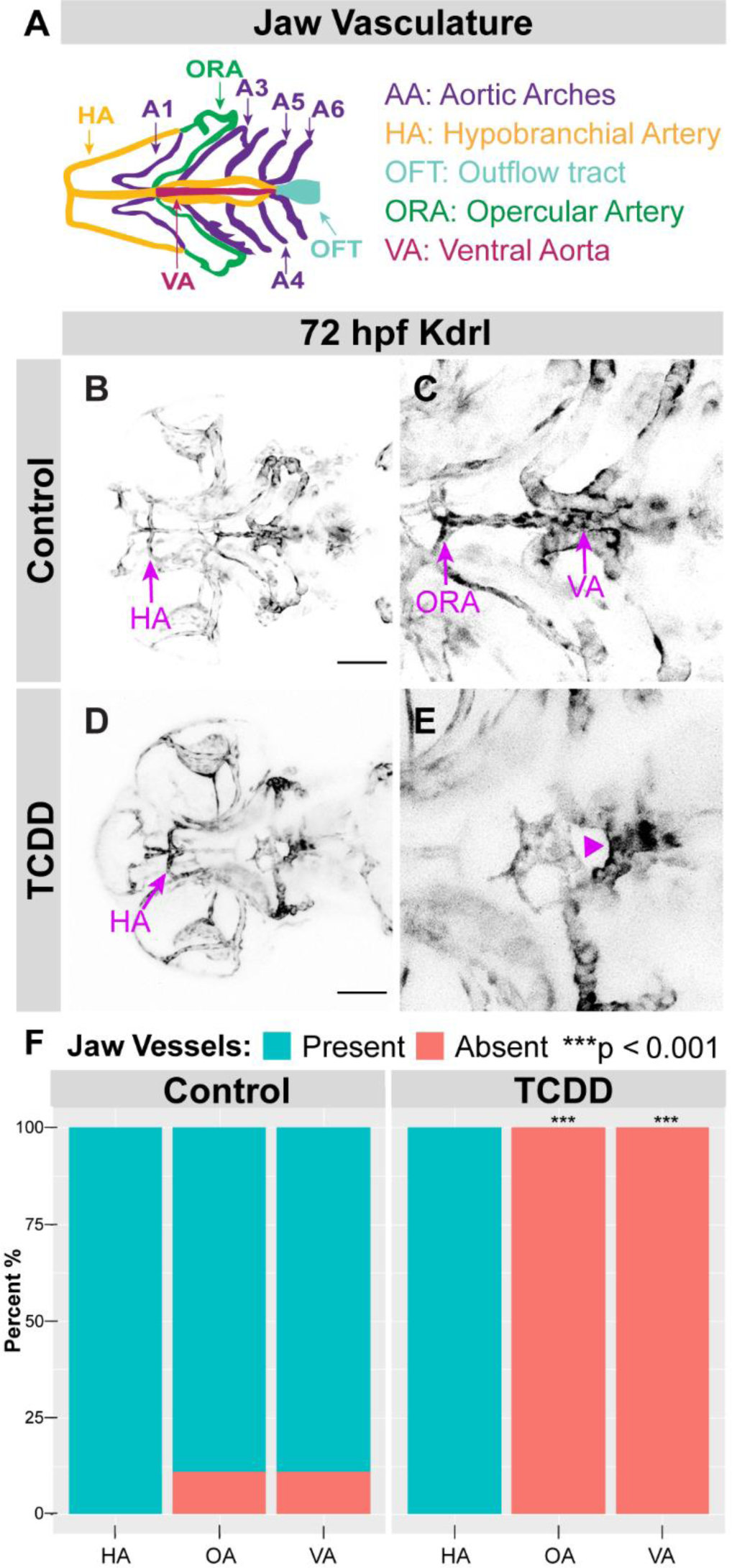
TCDD exposure leads to vasculature malformations. Schematic representations of developed zebrafish jaw vasculature from a ventral view (A). Maximum intensity projections of confocal micrographs at 10x magnification (B, D) and with a 1.5 digital zoom (C, E) capturing control (B, C) and TCDD-exposed (D, E) larvae at 72 hpf expressing reporters for *kdrl* (black) [*Tg(kdrl:GFP)*]. Stacked bar graphs indicating absence or presence of key jaw vessels (F). Arrowhead highlights absence of ventral aorta (E). Scale bar in B & D = 100μm. Total n in 72 hpf control group=9 and n in the TCDD-exposed group=9. Larvae were collected across 3 replicates. Abbreviations: hypobranchial artery (HA), opercular artery (OA), ventral aorta (VA). ***p < 0.001.

Our findings align with previous characterizations of the *tcf21* zebrafish mutant which showed that loss of Tcf21 resulted in severe reduction of the muscles of the head, but only mildly affected vasculature (Nagelberg et al., 2015). This previous study had indicated that in *tcf21* mutants all pharyngeal arteries developed properly and abnormalities were only detected in the hypobranchial artery shape and its connection to other vessels (Nagelberg et al., 2015). These milder vasculature phenotypes, compared to the severe absence of muscle phenotype, are likely due to the cross regulation of vasculature development orchestrated by Tcf21 in conjunction with Nkx2.5 (Nagelberg et al., 2015). Given its involvement in regulating vasculature development, Nkx2.5 should be further interrogated in future TCDD studies. Additionally, it is worth noting that the *tcf21* mutant fish only showed mild defects in cartilage formation as visualized through Alcian blue staining (Nagelberg et al., 2015), suggesting that loss of Tcf21+ progenitors alone is not responsible for the cartilage defects seen in this study following embryonic TCDD exposure. In brief, our data indicate that embryonic TCDD exposure depleted Sox10+ chondrocytes and Tcf21+ progenitors in the jaw and, ultimately, resulted in craniofacial cartilage, collagen, muscle, nerve, and vascular abnormalities (Fig 8).

**Figure 8.**
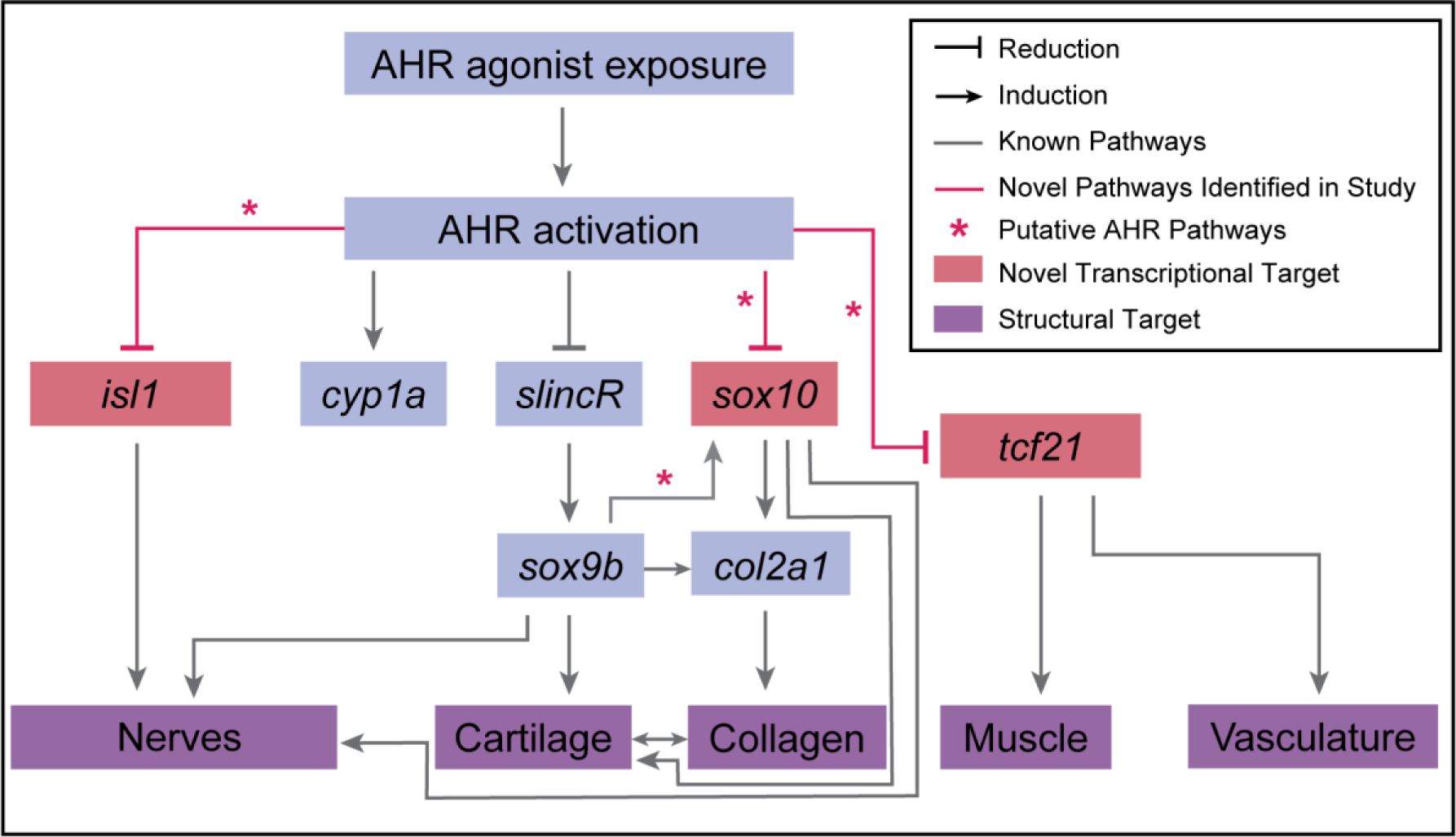
Summary of established and novel pathways mediating TCDD induced craniofacial toxicity. Schematic representation of transcriptional and structural targets of TCDD-induced toxicity. Asterisks indicate potential AHR pathways that require further investigation.

## 5 Conclusions

This study unveils novel targets of embryonic TCDD exposure and AHR-induced toxicity. Specifically, this work reveals Sox10+ chondrocytes as a cranial neural crest derived target and Tcf21+ progenitors as a novel pharyngeal mesoderm derived target. Through this work, we not only quantified the effects of TCDD exposure on expression of novel transcriptional targets, Sox10 and Tcf21, but also characterized the phenotypic effects induced by TCDD on critical craniofacial structures derived from or guided by Sox10+ neural crest cells and Tcf21+ progenitors. Our work indicates that embryonic exposure to TCDD results in reduction of Sox10+ chondrocytes and its associated collagen type II deposition. Likewise, TCDD exposed fish showed reduction in jaw and pharyngeal Tcf21+ progenitors which are required for axon guidance, muscle establishment, and vasculature development. Accordingly, TCDD exposed fish displayed impaired muscle formation, diminished overall ventral axon establishment, compression of cranial nerves V and VII, and vasculature malformations. Together our data demonstrate that embryonic exposure to TCDD targets multiple, previously undescribed, progenitors, cell types, and structures. Future work should focus on determining the hierarchy by which the early targets identified in this study induce the observed jaw and neurocranium phenotypes.

## 7 Author Contributions

L.G.C-R and J.S.P. designed the research study, developed figures, and wrote the manuscript. L.G.C-R performed TCDD exposures, collected, and prepared samples. L.G.C-R. and R.P. performed confocal imaging and acquired data for analysis. L.G.C-R and N.B. conducted data analysis. L.G.C-R is supported by the National Science Foundation Graduate Research Fellowship (NSF-GRFP and previously by the NIEHS Training in Environmental Pathology T32 (ES007272). J.S.P. is supported by a CPVB Phase II COBRE (2PG20GM103652) and an NIEHS ONES award (ES030109).

## 8 Acknowledgements

The authors would like to express their gratitude for all the past and present members of the Plavicki Lab. They would like to thank all members for their help in maintaining the lab and for their feedback throughout this project.

