## Supplemental Figures for "Exposure to the aryl hydrocarbon receptor agonist dioxin disrupts formation of the muscle, nerves, and vasculature in the developing jaw"

### 7 Supplemental Figures

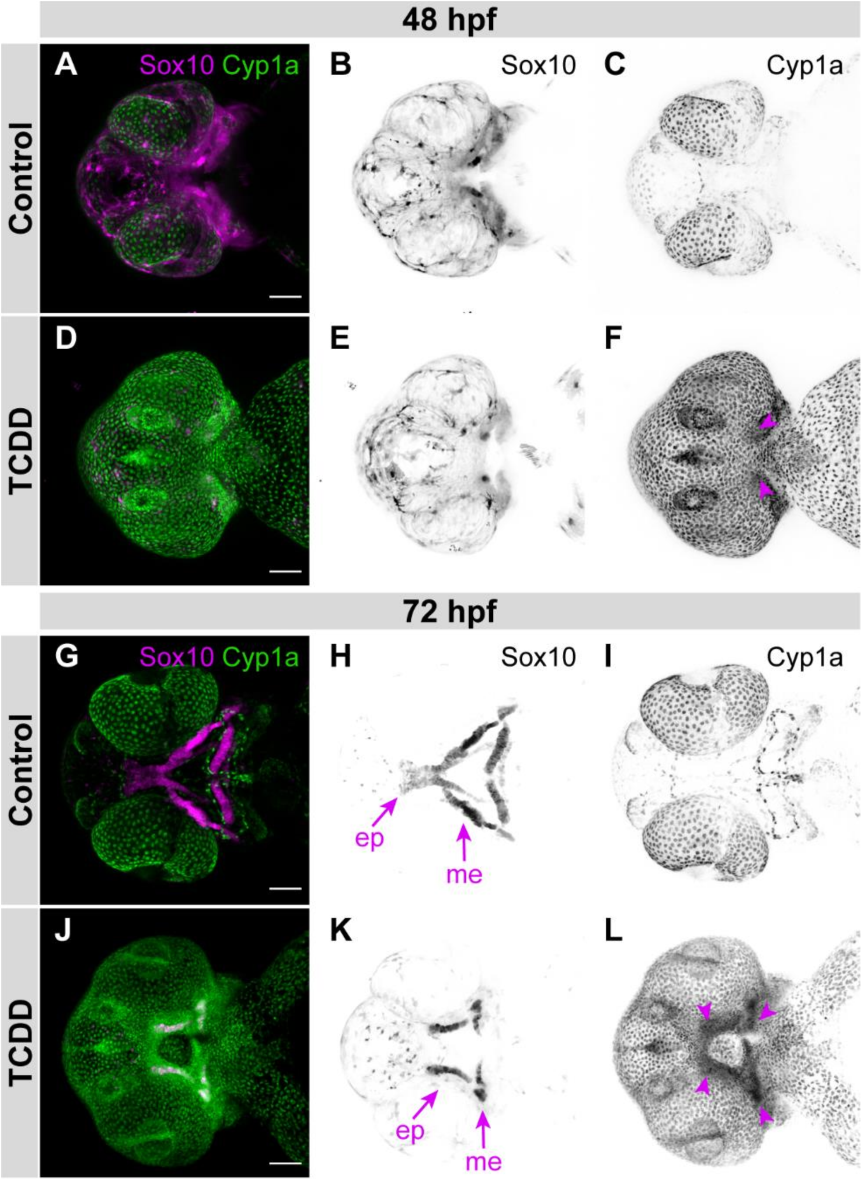


**Supplemental Figure 1. Exposure to TCDD induces *cyp1a* in the jaw.** Maximum intensity projections of confocal micrographs at 10x magnification capturing control (A-C, G-H) and TCDD-exposed (D-F, J-L) larvae at 48 hpf (A-F) and 72 hpf (G-L) expressing reporters for *sox10* (magenta) [*Tg(sox10:RFP)*] and *cyp1a* [*Tg (cyp1a:NLS:EGFP)*]. Maximum intensity projections of confocal z-series in black and white showing distinct expression patterns of Sox10 (B, E, H, & K) and Cyp1a (C, F, L), & I. Arrowheads indicate Cyp1a expression in Sox10+ domain (F, L). Scale bar in A, D, G, & J = 100μm. Abbreviations: Meckel's (me), ethmoid plate (ep).

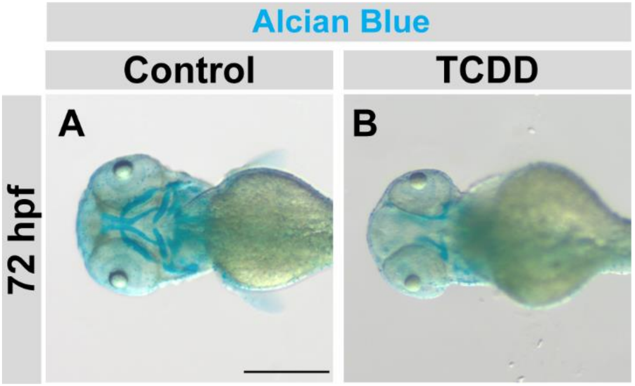


**Supplemental Figure 2. TCDD exposure reduced cartilage in the developing jaw.** Control (A) and TCDD (B) exposed larvae at 72 hpf stained for Alcian blue. Scale bar in A = 500 pixels. Total n in control group=9 and n in the TCDD-exposed group=9. Larvae were collected across 3 replicates.

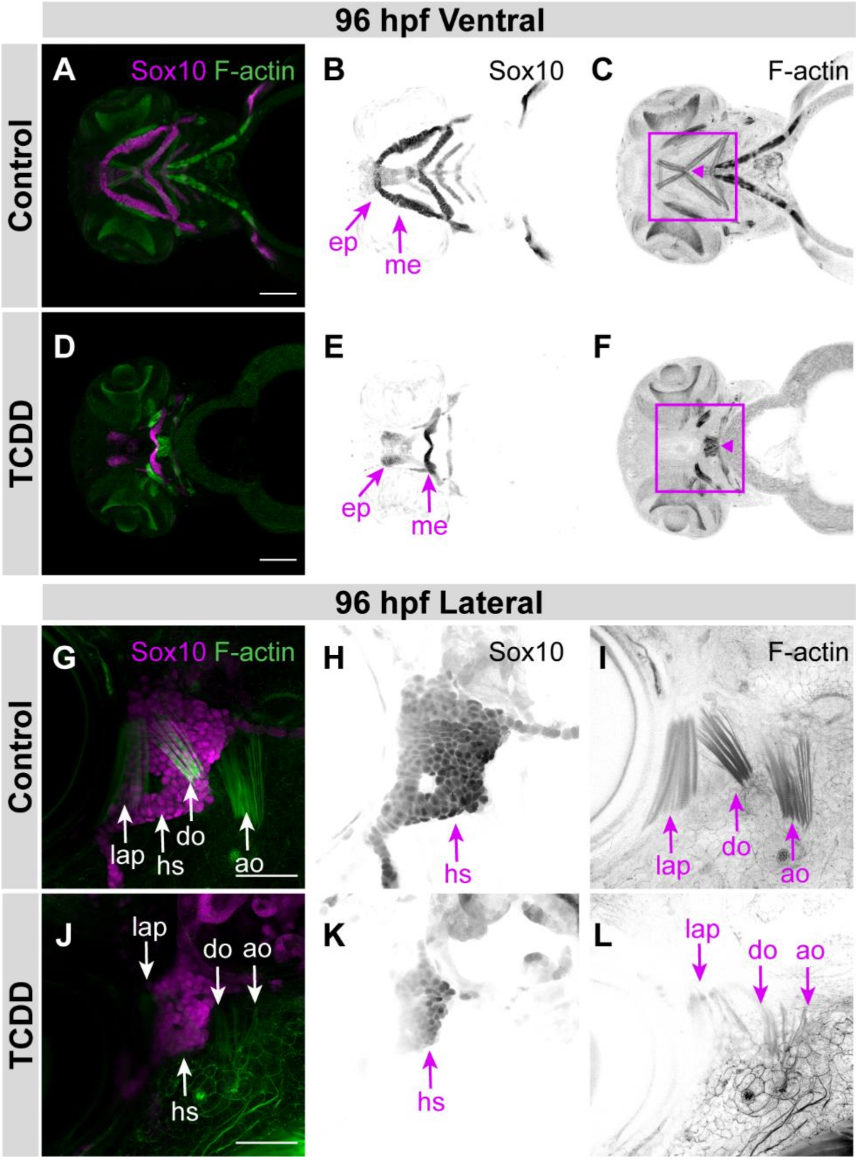


**Supplemental Figure 3. Representative images of Sox10+ chondrocytes and established muscles at 96 hpf.** Maximum intensity projections of confocal micrographs at 10x magnification capturing ventral 96 hpf control (A-C) and TCDD-exposed (D-F) larvae expressing reporters for *sox10* (magenta) [*Tg(sox10:RFP)*] and phalloidin stained (F-actin, green). Scale bar in A & D = 100μm. Confocal micrographs at 40x magnification capturing lateral 96 hpf control (G-I) and TCDD-exposed (J-L) larvae expressing reporters for *sox10* (magenta) [*Tg(sox10:RFP)*] and F-actin stained (green). Maximum intensity projections of confocal z-series in black and white showing distinct expression patterns of Sox10 (B, E, H, & K) and F-actin (C, F, I, & L). Box indicates jaw muscle domain and red arrowhead points to the junction between intermandibularis anterior and intermandibularis posterior (C, F). Scale bar in A & D = 100μm. Scale bar in G & J = 50μm. Abbreviations: hyosymplectic (hs), levator arcus palatini (lap), dilator operculi (do), and abductor operculi (ao).

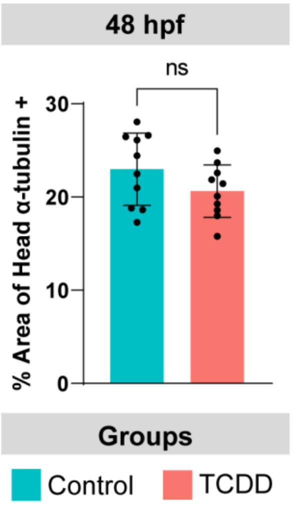


**Supplemental Figure 4. Ventral innervation at 48 hpf.** Bar graph of percentage area of ventral head covered by acetylated-α-tubulin expression at 48 hpf. Total n control group=13 and n in the TCDD-exposed group=12. Embryos were collected across 3 replicates. ns: $p>0.05$

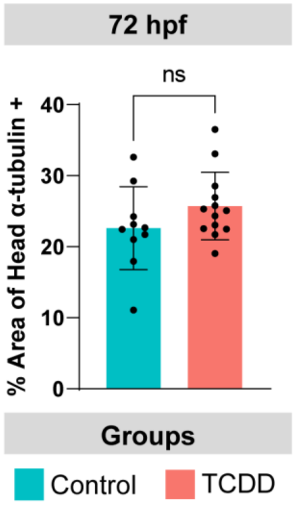


**Supplemental Figure 5. Lateral innervation at 72 hpf.** Bar graph of percentage area of lateral head covered by acetylated-α-tubulin expression at 72 hpf. Total n control group=24 and n in the TCDD-exposed group=23. Larvae were collected across 3 replicates. ns: $p>0.05$

**Supplemental Movie 1. Time-lapse microscopy of Sox10+ cells establishing cartilaginous jaw structures.** Control (Video 1) and TCDD exposed (Video 2) jaw development shown *in vivo* from 53 hpf – 70 hpf through confocal microscopy at 10x magnification. Total n in control group=2 and n in the TCDD-exposed group=2. Larvae were imaged across 2 replicates. Abbreviations: ethmoid plate (ep).

**Supplemental Movie 2. Time-lapse microscopy of Tcf21+ cells migrating to the jaw muscle domain.** Control (Video 1) and TCDD exposed (Video 2) jaw development shown in vivo from 53 hpf – 70 hpf through confocal microscopy at 10x magnification. Total n in control group=2 and n in the TCDD-exposed group=2. Larvae were imaged across 2 replicates. Abbreviations: intermandibularis anterior (ima), abductor mandibularis (am), hyohyal (hh), mandibulohyoid junction (mhj).
